# Associations Between Age, Heart Rate Variability, and BOLD fMRI Signal Variability

**DOI:** 10.1101/2025.11.26.690332

**Authors:** Jonathan Morris, Stacey M. Schaefer, Yiyi Zhu, Jinx Recchio, Lauren Gresham, Sarah E. Skinner, Kareem Al-Khalil, B. Suresh Krishna

## Abstract

Numerous studies report that BOLD fMRI signal variance (SD_BOLD_) decreases with age. However, these associations may partly reflect cardiovascular contributions to the BOLD signal. For example, heart rate variability (HRV) has been positively associated with Resting State Fluctuation Amplitude (RSFA), which captures low frequency components of BOLD fMRI variability. HRV is also negatively associated with age, which could potentially confound age-SD_BOLD_ associations. Yet, limited research has examined HRV-SD_BOLD_ associations or tested within-person HRV-SD_BOLD_ coupling using sliding window analyses of simultaneous HRV and SD_BOLD_. We analyzed resting-state fMRI data from two independent Midlife in the United States (MIDUS) samples: Core at M3 (n=115) and Refresher at MR1 (n=101). Partial Least Squares (PLS) analyses revealed significant positive HRV-SD_BOLD_ associations (Core: permutation p=0.018; Refresher: permutation p<0.001). Whole brain age-SD_BOLD_ PLS associations were non-significant via permutation tests across several models (Core: permutation p=0.201; Refresher: permutation p=0.121). We found age-related decreases in SD_BOLD_ across ∼70% of voxels in both samples. Concordance analyses showed 67-69% of brain voxels exhibited negative age-SD_BOLD_ but positive HRV-SD_BOLD_ relationships, suggesting that regions showing age-related decreases in SD_BOLD_ also showed HRV-related increases in SD_BOLD_. Sliding-window analyses demonstrated robust positive within-person associations between person-centered HRV and SD_BOLD_ via different HRV metrics: SDNN (Core: p < 0.001; Refresher: p < 0.001), RMSSD (Core: p = 0.072; Refresher: p = 0.009), and low frequency (Core: p < 0.001; Refresher: p < 0.001), with non-significant effects of high frequency (Core: p = 0.516; Refresher: p = 0.12) HRV. Thus, regardless of baseline levels, windows with higher HRV corresponded to higher SD_BOLD_, suggesting that cardiovascular factors partially explain age-SD_BOLD_ associations and HRV may mechanistically influence SD_BOLD_. These results suggest that controlling for HRV, especially low-frequency HRV or SDNN, may be necessary when analyzing SD_BOLD_ to isolate neural effects.

## Introduction

BOLD signal variability has emerged as an important metric for understanding brain function, reflecting the brain’s capacity for flexible information processing and adaptive responses to environmental demands (Garrett et al., 2010; McIntosh et al., 2008). Greater BOLD variability potentially reflects a more dynamic neural system capable of adapting to varying cognitive demands (Garrett et al., 2010; McIntosh et al., 2008). While BOLD variability has been associated with numerous behavioral and demographic factors (Steinberg & King, 2024), neural components of the BOLD signal are often entangled with physiological noise (Tsvetanov et al., 2021), leaving unclear whether behavioral associations with BOLD variance can be attributed to neural signals or cardiovascular noise (Logothetis, 2008; Tsvetanov et al., 2015).

Two important methodological approaches to measure voxel-wise BOLD variability include (1) the SD_BOLD_ method introduced by Garrett et al. (2010) and (2) the resting state fluctuation amplitude (RSFA) method from Kannurpatti & Biswal (2008). SD_BOLD_ quantifies variability by first dividing the BOLD time series into temporal blocks (∼10 fMRI volumes per block), scaling each block to a common mean intensity to control for scanner drift, then removing the mean signal within each block before calculating the standard deviation of the resulting time series (Garrett et al., 2010). This approach captures neural signal fluctuations across a broad frequency range while minimizing the influence of slow signal drift. In contrast, RSFA (analogous to ALFF; Zang et al., 2007) measures the standard deviation of band-pass filtered resting-state BOLD signals (0.01-0.1 Hz), emphasizing slower oscillations that are strongly influenced by vascular and physiological processes (Kannurpatti & Biswal, 2008). While both methods quantify temporal variability in the BOLD signal, they capture slightly different aspects of BOLD variability. Researchers claim that SD_BOLD_ more directly indexes neural signal dynamics, whereas RSFA may serve as a proxy for vascular contributions to the BOLD response (Garrett et al., 2010; Kannurpatti & Biswal, 2008; Tsvetanov et al., 2021).

Previous literature has exhibited robust negative associations between SD_BOLD_ and age (Garrett et al., 2010, 2013, 2017; Grady & Garrett, 2018; Guitart-Masip et al., 2016; Millar et al., 2020; Rieck et al., 2017). For instance, Garrett et al. (2010) found that younger adults (mean age = 25.79 ± 3.28) exhibited significantly higher SD_BOLD_ during fixation blocks between task-based fMRI scans when compared to older adults (mean age = 66.46 ± 8.25). In a follow-up study with the same participants, Garrett et al. (2013) found that younger adults also showed higher SD_BOLD_ across task states. Similarly, Guitart-Masip et al. (2016) found higher SD_BOLD_ in younger adults (mean age = 25.16 ± 2.27) vs. older adults (mean age = 70.33 ± 3.2), with age differences mediated by dopaminergic neurotransmission availability measured via PET scans. Additionally, Grady & Garrett (2018) examined differences in SD_BOLD_ during resting state fMRI scans, finding once again that younger adults (mean age = 23.7 ± 3.1 yrs) exhibited higher SD_BOLD_ than older adults (mean age = 71.1 ± 5.1 yrs). Negative associations between age and SD_BOLD_ are well documented in task and resting state fMRI studies. Based on these findings, researchers have concluded that greater SD_BOLD_ may indicate a more flexible neural system, and that this flexibility may decrease with increasing age (Garrett et al., 2010, 2017).

RSFA is also negatively associated with age, suggesting that the effects of aging on BOLD variability may be explained by cardiovascular and cerebrovascular factors (Tsvetanov et al., 2015, 2020, 2021). The BOLD signal is known to be a complex convolution of both vascular and neural signals (Logothetis, 2008). Tsvetanov et al. (2015) found that age-related differences in RSFA were statistically mediated by cardiovascular health measures including heart rate variability (HRV), but importantly not by neural variability measured via magnetoencephalography (MEG). Additional studies show age effects on voxel-wise RSFA become non-significant after regressing out cardiovascular components (Tsvetanov et al., 2021). However, these findings have been examined solely using RSFA as a measure of BOLD variability, which tends to be more susceptible to the influence of cardiovascular noise (Tsvetanov et al., 2015). Garrett et al. (2017) argued against these vascular confound explanations, showing that age-related differences in SD_BOLD_ remained robust even after controlling for multiple vascular measures including absolute cerebral blood flow and BOLD cerebrovascular reactivity (CVR). BOLD CVR represents the capacity of cerebral blood vessels to dilate or constrict in response to vasoactive stimuli, typically measured by quantifying changes in the BOLD signal per unit change in end-tidal CO₂ during hypercapnia challenges such as breath-holding tasks or CO₂ inhalation (Garrett et al., 2017). However, Garrett et al. did not directly adjust for HRV.

HRV appears to be negatively associated with age up until around age fifty (Antelmi et al., 2004; Tegegne et al., 2018; Voss et al., 2015; Zhang, 2007), whereafter decreases in HRV plateau or even increase as shown in Antelmi et al. (2004). This decline is attributed to various age-related changes in health and autonomic nervous system function, including arterial stiffening, blood pressure changes, reduced parasympathetic activity, and increased sympathetic dominance (Tegegne et al., 2018). These reductions in HRV with age, paired with negative associations between BOLD variance and age suggest that age-related reductions in BOLD variance may at least partly be due to cardiovascular components such as HRV.

While past research from Tsvetanov et al. (2021) indicates that the effect of age on RSFA is dependent on vascular factors including HRV, no research has examined direct associations between HRV and BOLD variance directly using SD_BOLD_. SD_BOLD_ includes higher-frequency components that are filtered out of the RSFA and is considered to be more neural than RSFA; direct associations between HRV and SD_BOLD_ could suggest that broader BOLD frequency ranges may be susceptible to cardiovascular confounding. The present research provides the first direct test of associations between HRV and SD_BOLD_, further examining whether cardiovascular dynamics underlie age-related reductions in BOLD signal variance.

Using data from the Midlife in the United States (MIDUS) study featuring large age ranges spanning over 7 decades, we examined two separate datasets (MIDUS Core and MIDUS Refresher) to test associations between HRV, age, and SD_BOLD_ during resting state fMRI scans. This paper makes three key methodological contributions to understanding cardiovascular associations with BOLD signal variability. We measured HRV simultaneously during resting-state fMRI acquisition using photoplethysmography, whereas previous papers used cardiovascular measures obtained during separate sessions (Garrett et al., 2017; Tsvetanov et al., 2021). Additionally, we employed Partial Least Squares (PLS) analyses to examine HRV-SD_BOLD_ associations and identify distributed spatial patterns of brain activity associated with cardiovascular changes. Finally, we conducted novel within-person sliding window analyses to test whether moment-to-moment fluctuations in HRV predict concurrent changes in SD_BOLD_ within individuals, distinguishing true dynamic coupling from confounding by stable between-person differences. Robust positive associations between HRV and SD_BOLD_ could imply that negative associations between age and SD_BOLD_ may be driven by mechanistic cardiovascular effects on BOLD variability, emphasizing the need to properly control for cardiovascular factors when analyzing resting state fMRI data.

We initially examined direct associations between age and HRV. Second, using PLS, we separately tested associations between age, HRV, and SD_BOLD_. We also examined joint HRV and age models, testing whether HRV or age explained more variance in SD_BOLD_. We predicted that age would be negatively associated with both HRV and SD_BOLD_. Additionally, we hypothesized positive associations between HRV and SD_BOLD_ and that in joint PLS models, HRV would explain more variance in SD_BOLD_ than age. Lastly, using sliding window analyses to test within-person coupling between HRV and average SD_BOLD_, we predicted that windows with higher HRV would also have higher average SD_BOLD_ regardless of individual baseline HRV.

## Method

### Study Design and Participants

Data were analyzed from two independent datasets from the Midlife in the United States (MIDUS) study: Core sample at MIDUS 3 (M3; n=115; age: M=65.9 years, SD=9.1, range=48-95; 42% male; 80% White, 20% Black, Indigenous, and people of color (BIPOC)) and Refresher sample at MIDUS Refresher 1 (MR1; n=102; age: M=48.5 years, SD=11.6, range=26-76; 53% female; 69% White, 31% BIPOC). MIDUS 3 neuroscience data were collected between 2017-2022 from participants in the original longitudinal MIDUS Core cohort (first recruited in 1995-1996). MIDUS Refresher 1 neuroscience data were collected between 2012-2015 from a newly recruited national probability sample (MIDUS Refresher, enrolled 2011-2014) designed to provide age-matched cross-sectional comparisons to the original MIDUS Core cohort. Both studies included participants who completed task free (resting-state) functional magnetic resonance imaging (fMRI) scans and had complete physiological heart rate variability (HRV) data acquired simultaneously through photoplethysmography (PPG).

### MRI Data Acquisition

#### Imaging Data Acquisition

MRI data were collected on the same 3T scanner (MR750 GE Healthcare, Waukesha, WI) using an 8-channel head coil for the Refresher sample at MIDUS Refresher 1 and a 32-channel head coil for the Core sample at MIDUS 3. Structural T1-weighted images were acquired using a 3-dimensional magnetization-prepared rapid gradient-echo sequence with the following parameters: TR=8.2ms, TE=3.2ms, flip angle=12°, field of view=256mm, 256×256 matrix, 160 axial slices, inversion time=450ms. Resting-state functional MRI data were acquired using echo planar imaging with the following parameters: 240 volumes, TR=2000ms, TE=20ms, flip angle=60°, field of view=220mm, 96×64 matrix, 3-mm slice thickness with 1-mm gap, 40 interleaved sagittal slices, and ASSET parallel imaging with an acceleration factor of 2. During resting-state scans, participants were instructed to lie still with their eyes open.

### Physiological Data Collection and Processing

Psychophysiology data (skin conductance response, respiration, photoplethysmography) were gathered during fMRI scans in both samples using BIOPAC systems at 1000Hz. In the Refresher sample, an MP150 system was used throughout. In the Core sample, an MP150 system was initially used but was replaced with an MP160 system mid-study (August 2021), with some participants having pulse oximetry collected on different amplifiers (OXY100C vs PPG100C for mp150 and mp160, respectively). HRV measures were derived from the photoplethysmography (PPG) data collected during resting-state fMRI acquisition.

Using NeuroKit2 version 0.2.11 (Makowski et al., 2021), heart rate variability (HRV) measures were extracted from photoplethysmography (PPG) data collected during resting-state scanning sessions. PPG signals were cleaned using NeuroKit2’s standard cleaning algorithm and peaks were identified with artifact correction enabled. Manual quality control was performed through an interactive visual inspection process where a trained rater identified and marked segments containing artifacts or signal dropout. These marked segments were converted to NaN values in the peak timestamp series, effectively excluding them from HRV calculations while preserving the overall temporal structure of the data. Time-domain measures included root mean square of successive differences (RMSSD) and standard deviation of normal-to-normal intervals (SDNN). Frequency-domain measures were calculated using Fast Fourier Transform (FFT) with an interpolation rate of 10 Hz, yielding low-frequency power (LF; 0.04-0.15 Hz) and high-frequency power (HF; 0.15-0.40 Hz). To account for skewness in the HRV data distribution, log-transformed measures were used in all analyses.

### fMRI Preprocessing

Our preprocessing pipeline followed all preprocessing steps conducted by Garrett et al. (2010). Results included in this manuscript come from preprocessing performed using *fMRIPrep* 23.0.0 (Esteban, Markiewicz, et al., (2018); Esteban, Blair, et al., (2018); RRID:SCR_016216), which is based on *Nipype* 1.8.5 (K. Gorgolewski et al., (2011); K. J. Gorgolewski et al., (2018); RRID:SCR_002502). The following preprocessing description was pasted from the fMRIPrep-generated methods boilerplate.

#### Anatomical Data Preprocessing

For each subject the following anatomical preprocessing was conducted. The T1-weighted (T1w) image was corrected for intensity non-uniformity (INU) with N4BiasFieldCorrection (Tustison et al., 2010), distributed with ANTs 2.3.3 (Avants et al., 2008; RRID:SCR_004757), and used as T1w-reference throughout the workflow. The T1w-reference was then skull-stripped with an *Nipype* implementation of the antsBrainExtraction.sh workflow (from ANTs), using OASIS30ANTs as target template. Brain tissue segmentation of cerebrospinal fluid (CSF), white-matter (WM) and gray-matter (GM) was performed on the brain-extracted T1w using fast (FSL 6.0.5.1:57b01774, RRID:SCR_002823, Zhang, Brady, and Smith, 2001). Volume-based spatial normalization to two standard spaces (MNI152NLin2009cAsym, MNI152NLin6Asym) was performed through nonlinear registration with antsRegistration (ANTs 2.3.3), using brain-extracted versions of both T1w reference and the T1w template. The following templates were selected for spatial normalization: *ICBM 152 Nonlinear Asymmetrical template version 2009c* (Fonov et al., 2009; RRID:SCR_008796; TemplateFlow ID: MNI152NLin2009cAsym), *FSL’s MNI ICBM 152 non-linear 6th Generation Asymmetric Average Brain Stereotaxic Registration Model* (Evans et al., 2012; RRID:SCR_002823; TemplateFlow ID: MNI152NLin6Asym).

#### Functional data preprocessing

Each subject’s BOLD run was pre-processed as follows. First, a reference volume and its skull-stripped version were generated using *fMRIPrep*. Head-motion parameters with respect to the BOLD reference (transformation matrices, and six corresponding rotation and translation parameters) were estimated before any spatiotemporal filtering using mcflirt (FSL 6.0.5.1:57b01774; Jenkinson et al., 2002). BOLD runs were slice-time corrected to 0.975s (0.5 of slice acquisition range 0s-1.95s) using 3dTshift from AFNI (Cox and Hyde, 1997; RRID:SCR_005927). The BOLD time-series (including slice-timing correction when applied) were resampled onto their original, native space by applying the transforms to correct for head-motion. These resampled BOLD time-series will be referred to as *preprocessed BOLD in original space*, or just *preprocessed BOLD*. The BOLD reference was then co-registered to the T1w reference using mri_coreg (FreeSurfer) followed by flirt (FSL 6.0.5.1:57b01774; Jenkinson and Smith, 2001) with the boundary-based registration (Greve and Fischl, 2009) cost-function. Co-registration was configured with six degrees of freedom. Several confounding time-series were calculated based on the *preprocessed BOLD*: framewise displacement (FD), DVARS and three region-wise global signals. FD was computed using two formulations following Power (absolute sum of relative motions, Power et al., 2014) and Jenkinson (relative root mean square displacement between affines, Jenkinson et al., 2002). FD and DVARS were calculated for each functional run, both using their implementations in *Nipype* (following the definitions by Power et al., 2014). The three global signals were extracted within the CSF, the WM, and the whole-brain masks. Additionally, a set of physiological regressors were extracted to allow for component-based noise correction (*CompCor*; Behzadi et al., 2007). Principal components were estimated after high-pass filtering the *preprocessed BOLD* time-series (using a discrete cosine filter with 128s cut-off) for the two *CompCor*variants: temporal (tCompCor) and anatomical (aCompCor). tCompCor components were then calculated from the top 2% variable voxels within the brain mask. For aCompCor, three probabilistic masks (CSF, WM and combined CSF+WM) were generated in anatomical space. The implementation differs from that of Behzadi et al. in that instead of eroding the masks by 2 pixels on BOLD space, a mask of pixels that likely contain a volume fraction of GM was subtracted from the aCompCor masks. This mask was obtained by thresholding the corresponding partial volume map at 0.05, which ensured components were not extracted from voxels containing a minimal fraction of GM. Finally, these masks were resampled into BOLD space and binarized by thresholding at 0.99 (as in the original implementation).Components were also calculated separately within the WM and CSF masks. For each CompCor decomposition, the *k* components with the largest singular values were retained, such that the retained components’ time series were sufficient to explain 50 percent of variance across the nuisance mask (CSF, WM, combined, or temporal). The remaining components were dropped from consideration. The head-motion estimates calculated in the correction step were also placed within the corresponding confounds file. The confound time series derived from head motion estimates and global signals were expanded with the inclusion of temporal derivatives and quadratic terms for each (Satterthwaite et al., 2013). Frames that exceeded a threshold of 0.5 mm FD or 1.5 standardized DVARS were annotated as motion outliers. Additional nuisance timeseries were calculated by means of principal components analysis of the signal found within a thin band (*crown*) of voxels around the edge of the brain, as proposed by (Patriat, Reynolds, and Birn, 2017).

The BOLD time-series were resampled into several standard spaces, correspondingly generating the following *spatially-normalized, preprocessed BOLD runs*: MNI152NLin2009cAsym, MNI152NLin6Asym. First, a reference volume and its skull-stripped version were generated. Automatic removal of motion artifacts using independent component analysis (ICA-AROMA; Pruim et al., 2015) was performed on the *preprocessed BOLD on MNI space*time-series after removal of non-steady state volumes and spatial smoothing with an isotropic, Gaussian kernel of 6mm FWHM (full-width half-maximum). Corresponding “non-aggresively” denoised runs were produced after such smoothing. Additionally, the “aggressive” noise-regressors were collected and placed in the corresponding confounds file. All resamplings were performed with *a single interpolation step* by composing all the pertinent transformations (i.e. head-motion transform matrices, susceptibility distortion correction when available, and co-registrations to anatomical and output spaces). Gridded (volumetric) resamplings were performed using antsApplyTransforms (ANTs), configured with Lanczos interpolation to minimize the smoothing effects of other kernels (Lanczos, 1964). Non-gridded (surface) resamplings were performed using mri_vol2surf (FreeSurfer). Following *fMRIPrep*, the first 5 volumes from each BOLD time series were removed to eliminate T1 saturation effects. Extended preprocessing steps included white matter (WM) and cerebrospinal fluid (CSF) signal extraction and regression using confounds created by *fMRIPrep*. BOLD data were spatially smoothed using an 8mm full-width half-maximum (FWHM) Gaussian kernel.

For both datasets, BOLD signal variability at each voxel was quantified following the SD_BOLD_ methodology established by Garrett et al. (2010). Voxel-wise BOLD time series were separated into temporal blocks (10 volumes per block). First, for each block separately, the overall 4D mean across all spatial dimensions and time was calculated. Each block was then scaled by multiplying all values by a normalization factor (100 / 4D mean), ensuring that the 4D mean of each block equaled 100. Second, for each voxel independently, the temporal mean within each block was calculated and subtracted from all time points in that block, centering each voxel’s time series around zero within each block. The mean-centered blocks were then concatenated across time for each voxel. The standard deviation at each voxel of the resulting concatenated time series was calculated to yield maps of SD_BOLD_ for each subject.

### Analyses

All analyses were conducted using RStudio, Python 3.10 with neuroimaging libraries nibabel (version 5.3.2), nilearn (version 0.12.1), and FSL (version 6.0.7.7).

#### Regression Analyses HRV-Age

We used simple linear regression to test the relationship between age and mean SD_BOLD_. Each HRV measure (HF, LF, RMSSD, and SDNN) was tested in a separate regression model to avoid collinearity effects. We applied Benjamini-Hochberg false discovery rate (FDR) correction to account for multiple comparisons. To control for sex and race, all linear models included sex (male vs female) and race (white vs BIPOC) as categorical factors.

#### Partial Least Squares (PLS)

PLS identifies multivariate spatial patterns of SD_BOLD_ that optimally relate to the external variables of interest (i.e., age and HRV measures). This approach is well-suited for neuroimaging applications because it captures distributed brain activity patterns rather than focusing on individual voxels in isolation. Three distinct PLS analyses were conducted identically across both MIDUS datasets: (1) age as sole predictor, (2) four HRV measures (SDNN, RMSSD, LF, HF) as predictors, and (3) joint modeling of age with all four HRV measures.

All HRV measures were included simultaneously in PLS analyses (2) and (3). PLS creates latent variables that maximize the covariance between the explanatory variables and SD_BOLD_. This method is robust to collinearity from multiple predictors, as the latent variables capture the covariance between multiple measures of HRV or age, capturing a combined effect of the respective variables. Prior to analysis, predictor variables and each voxel’s SD_BOLD_ values were z-scored across subjects (mean=0, SD=1) to place all variables on a common scale and facilitate directional interpretation of PLS results. Z-scoring predictor variables ensures that associations with brain scores can be interpreted relative to the sample mean, while z-scoring voxels standardizes variance magnitude across brain regions without removing meaningful between-subject differences in SD_BOLD_ patterns.

The PLS procedure constructs correlation matrices between each predictor variable and SD_BOLD_ values across all brain voxels computed across participants. These correlation matrices undergo singular value decomposition (SVD), which extracts latent variables characterized by two key components: *singular values* (reflecting correlation magnitude) and *voxel salience* patterns (representing optimal voxel weightings that capture relationships between SD_BOLD_ and explanatory variables, i.e., age and HRV). The voxel saliences function as unit-length vectors that identify which voxels most strongly contribute to the observed brain-behavior associations. The sign of these voxel saliences indicates whether typical multivariate patterns show increases or decreases in SD_BOLD_.

Individual participant *brain scores* were derived by computing the dot product between each subject’s whole-brain SD_BOLD_ pattern and the extracted voxel salience patterns. These brain scores quantify how strongly each participant expresses the identified multivariate spatial pattern. The interpretation of PLS results requires joint consideration of brain scores and voxel saliences: when participants with higher predictor values (e.g., older age) show elevated brain scores, and specific voxels exhibit negative saliences, this indicates that those brain regions demonstrate decreased SD_BOLD_ with increasing predictor values. Similarly, when participants with lower predictor values (e.g., lower HRV) exhibit higher brain scores, and voxels in a specific region have negative brain saliences, this suggests that those regions show positive correlations between the predictor and SD_BOLD_.

For age-only and HRV-only models, we focused our analyses on the first latent variable, as these represented either the dominant pattern of covariation between SD_BOLD_ and HRV (for the 4-variable HRV-only models), or the only relevant pattern (for the model with age as the only explanatory variable). However, for joint age and HRV models, we examined both the first and second latent variables to capture potentially divergent effects stemming from age and HRV. The proportion of positive versus negative voxel saliences, combined with brain score patterns, allows interpretation of whether SD_BOLD_ increases or decreases across distributed brain networks as a function of predictor variables.

Statistical significance was established through permutation testing (1000 iterations) applied to singular values, with alpha set at p < 0.05. To assess the spatial reliability of our findings, we implemented bootstrap resampling (1000 iterations) to generate bootstrap ratios for each voxel by dividing mean bootstrap saliences by their standard errors. Voxels surpassing a bootstrap ratio threshold of ±2.70 (approximating 99% confidence intervals) were considered spatially robust (Krishnan et al., 2011).

Concordance analyses were performed to examine spatial correspondence between age-only (analysis 1) and HRV-only (analysis 2) models. Each voxel was classified based on the directionality of effects: concordant positive (both age and HRV associated with increased SD_BOLD_), concordant negative (both associated with decreased SD_BOLD_), opposite directions (age and HRV showing opposing associations), or negligible effects. This analysis quantified the degree to which brain regions showing age-related changes in SD_BOLD_ overlapped with regions showing HRV-related changes.

#### Within-Person Analyses

In addition to voxel-wise PLS analyses, we conducted within-person sliding window analyses of the relationship between SD_BOLD_ and HRV. For both MIDUS Core and MIDUS Refresher, we divided the 8-minute resting state scan into 3 windows of 4 minutes each (50% overlap). Within each window, we calculated HRV and average SD_BOLD_ across all voxels for each subject using methods described above. Windows with signal dropout or excessive artifacts greater than two minutes were excluded from analysis. To isolate within-person changes in HRV, we calculated person-centered HRV using the following procedure. For each measure of HRV (SDNN, LF, HF, RMSSD), we computed subject-specific standard deviation and mean HRV across windows. Then for each subject HRV observation, we subtracted the subject-specific mean and divided by the subject-specific standard deviation.

Using the *lme4* package in RStudio, linear mixed-effects models tested whether within-person deviations in HRV predicted concurrent fluctuations in average SD_BOLD_. To ensure model convergence and avoid collinearity, we ran separate mixed effects models for each HRV variable and age. Each model included the person-centered and standardized HRV variable, which captured within-person deviations from an individual’s mean HRV, and the person-mean HRV variable, which measured between-person differences in average HRV across the scan. Age, sex, and race were included as covariates. Random intercepts and random slopes for within-person HRV predictors were estimated for each participant to account for individual differences in both baseline SD_BOLD_ and the strength of HRV–SD_BOLD_ coupling. Positive associations between person-centered HRV and SD_BOLD_ indicate that within participants, windows with higher HRV simultaneously have higher SD_BOLD_. Positive associations between person-mean HRV and SD_BOLD_ indicate that higher average HRV is associated with higher average SD_BOLD_ between participants. We predicted robust positive associations between both person-centered and person-mean HRV with SD_BOLD._

## Results

### Associations Between Age and HRV

In the Core sample, age was not negatively associated with HRV (**Fig. 1)**, with non-significant associations between age and high frequency HRV (β(114) = 0.82, p = .174), low frequency HRV (β(114) = -0.10, p = .879), and SDNN (β(114) = 2.20, p = .144), and a significant positive association between age and RMSSD (β(114) = 2.55, p = .016). The Refresher sample exhibited consistent age-related declines across multiple HRV indices, including high frequency HRV (β(101) = -2.14, p = .014), low frequency HRV (β(101) = -2.39, p = .0125), and SDNN (β(101) = -5.73, p = .014), with RMSSD showing a marginal negative association (β(101) = -3.32, p = .061). All HRV measures were tested in separate models to avoid collinearity. After applying Benjamini-Hochberg false discovery rate (FDR) correction for multiple comparisons within each sample, no associations between age and HRV in the Core sample remained significant, while three of four HRV indices in the Refresher sample (SDNN, high frequency, and low frequency HRV) retained significant associations.

**Figure 1:**
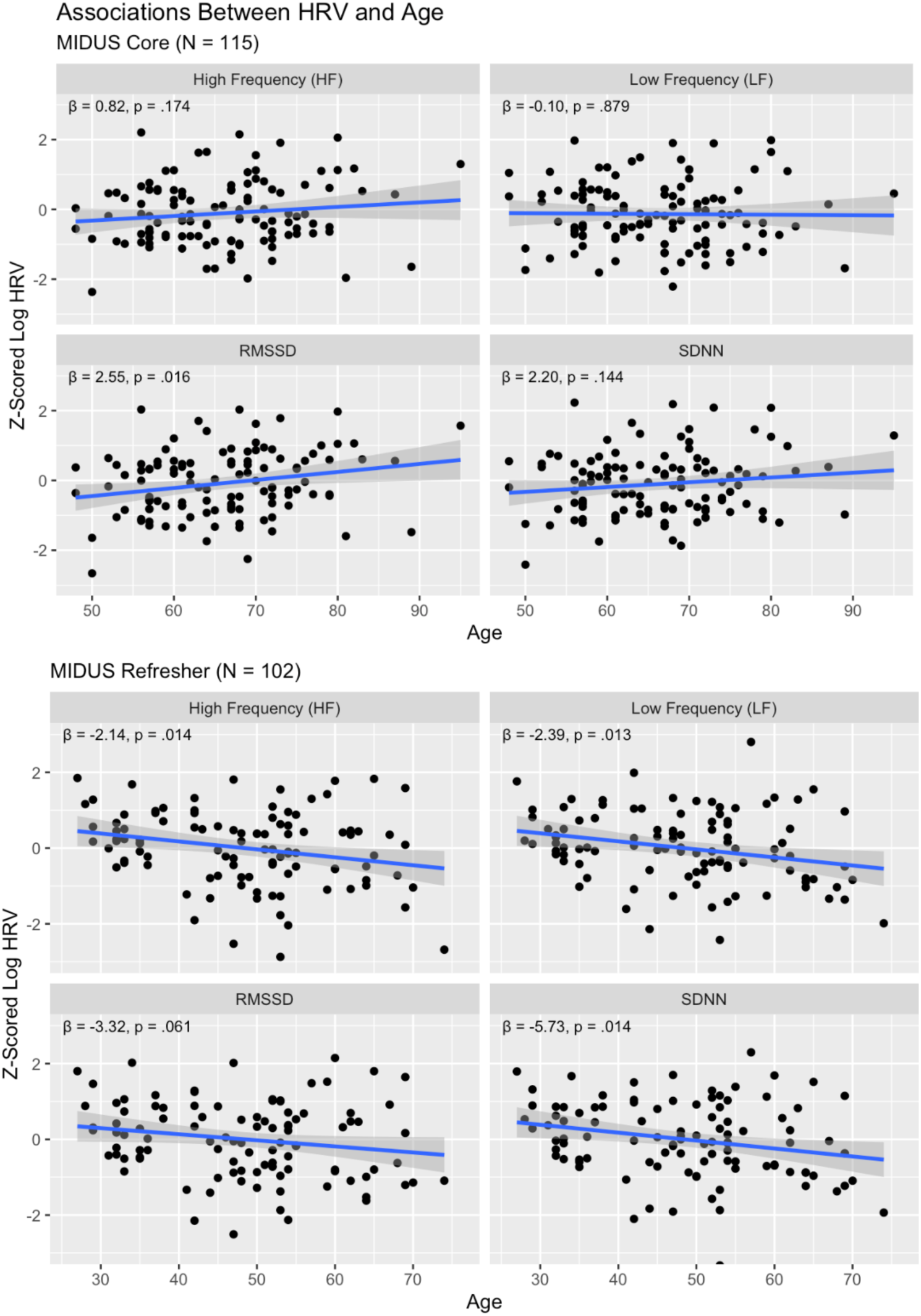
Associations Between HRV and Age. Scatterplots show associations between age and HRV across HRV measures and respective samples. For MIDUS Core, no significant associations between age and HRV were observed after false discovery rate (FDR) correction. For MIDUS Refresher, associations between age and three HRV indices (HF, LF, and SDNN) passed FDR correction, while RMSSD did not.

### Age-Related Differences in BOLD Signal Variability

*Core Results:* PLS analysis examining the relationship between age and SD_BOLD_ revealed a non-significant relationship (permutation p-value = 0.201), despite showing the expected directional pattern. Age was significantly associated with brain scores (β(114) = 5.54, t = 3.46, p < 0.001), with 70.1% of brain voxels showing negative voxel weights, indicating decreased variability with age across most brain regions (**Tables 1, 2**). Bootstrap significance testing revealed that 5.9% of voxels passed the stringent bootstrap ratio threshold of ±2.70, with 94% of voxels passing bootstrap ratio tests exhibiting negative voxel weights (**Fig. 2**).

**Figure 2:**
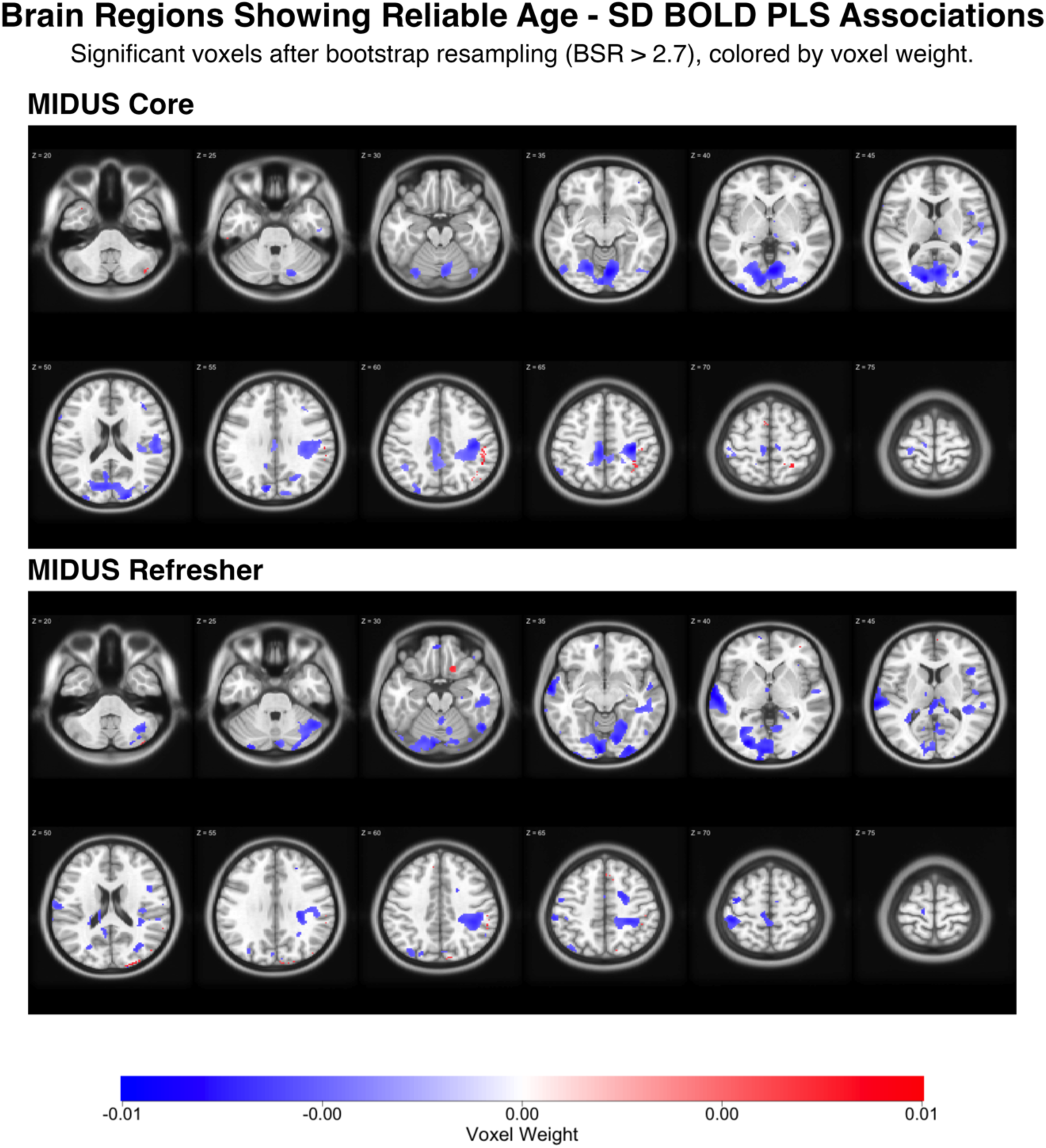
Age-Related Patterns in BOLD Signal Variability Across Core and Refresher Samples. Brain regions showing statistically reliable associations between age and BOLD signal variability (SD_BOLD_) derived from partial least squares (PLS) analysis. The top panel displays results from the MIDUS Core sample (n=115), while the bottom panel shows the MIDUS Refresher sample (n=102). Axial brain slices progress from inferior to superior, displaying voxels that passed bootstrap significance testing (bootstrap ratio > 2.7, approximately 99% confidence interval). Blue regions indicate negative associations between age and SD_BOLD_ (decreased variability with increasing age), while red regions indicate positive associations (increased variability with increasing age).

**Table 1.**
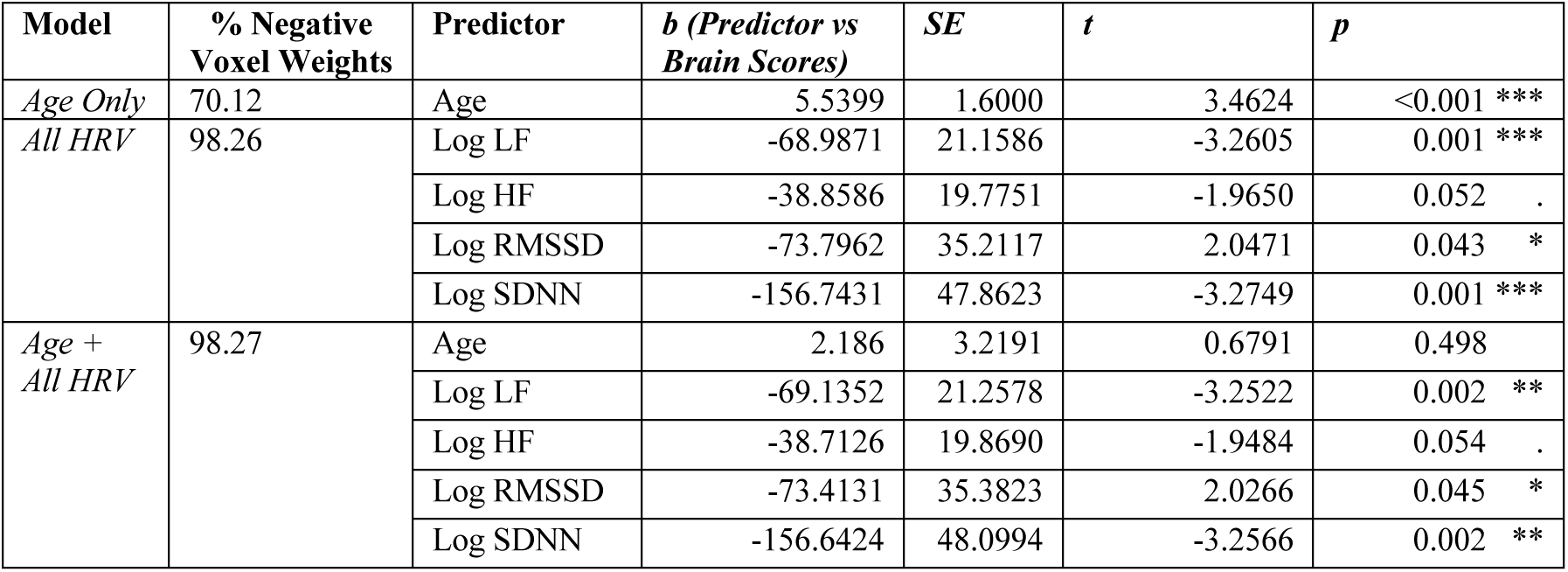
MIDUS Core PLS Results (Voxel Weights and Brain Scores)

| Model | % Negative Voxel Weights | Predictor | $b$ (Predictor vs Brain Scores) | SE | $t$ | $p$ |
| --- | --- | --- | --- | --- | --- | --- |
| Age Only | 70.12 | Age | 5.5399 | 1.6000 | 3.4624 | <0.001 *** |
| All HRV | 98.26 | Log LF | -68.9871 | 21.1586 | -3.2605 | 0.001 *** |
|  |  | Log HF | -38.8586 | 19.7751 | -1.9650 | 0.052 . |
|  |  | Log RMSSD | -73.7962 | 35.2117 | 2.0471 | 0.043 * |
|  |  | Log SDNN | -156.7431 | 47.8623 | -3.2749 | 0.001 *** |
| Age + All HRV | 98.27 | Age | 2.186 | 3.2191 | 0.6791 | 0.498 |
|  |  | Log LF | -69.1352 | 21.2578 | -3.2522 | 0.002 ** |
|  |  | Log HF | -38.7126 | 19.8690 | -1.9484 | 0.054 . |
|  |  | Log RMSSD | -73.4131 | 35.3823 | 2.0266 | 0.045 * |
|  |  | Log SDNN | -156.6424 | 48.0994 | -3.2566 | 0.002 ** |

**Table 2.** MIDUS Core PLS Permutation and Bootstrap Results.

| Model | Permutation P-Value | Bootstrap 95% CI | % Voxels Passing Bootstrap Ratio Test ($\pm 2.70$) |
| --- | --- | --- | --- |
| Age Only | 0.201 | [0.2618, 0.7185] | 5.9039 |
| All HRV | 0.018 ** | [0.2412, 0.4375] | 31.4964 |
| Age + All HRV | 0.019 ** | [0.2512, 0.4387] | 31.7961 |

#### Refresher Results

Similarly, the age-only PLS model showed a non-significant relationship (permutation p-value = 0.121), though with a stronger effect size than MIDUS Core. Age was significantly associated with brain scores (β(101) = 5.15, t = 3.71, p < 0.001), and 67.7% of brain voxels displayed negative weights (**Tables 3, 4**), suggesting decreased variability with age. Bootstrap testing indicated that 6.9% of voxels exceeded the bootstrap ratio threshold, suggesting slightly more extensive spatial coverage than MIDUS Core but still limited overall (**Fig. 2**). Further analyses revealed that 95% of voxels passing bootstrap ratio tests exhibited negative voxel weights.

**Table 3.** MIDUS Refresher PLS Results (Voxel Weights and Brain Scores)

| Model | % Negative Voxel Weights | Predictor | $\beta$ (Predictor vs Brain Scores) | SE | $t$ | $p$ |
| --- | --- | --- | --- | --- | --- | --- |
| Age Only | 67.698 | Age | 5.1394 | 1.3855 | 3.7093 | <0.001 *** |
| All HRV | 96.250 | Log LF | -123.8761 | 21.181 | -5.8485 | <0.001 *** |
|  |  | Log HF | -68.7631 | 21.1370 | -3.2431 | 0.002 ** |
|  |  | Log RMSSD | -143.9306 | 42.7953 | -3.3632 | 0.001 *** |
|  |  | Log SDNN | -257.2044 | 54.0102 | -4.7621 | <0.001 *** |
| Age + All HRV | 96.022 | Age | 3.1831 | 2.508 | 1.2692 | 0.207 |
|  |  | Log LF | -123.7266 | 21.1122 | -5.8604 | <0.001 *** |
|  |  | Log HF | -68.8285 | 21.2030 | -3.2563 | 0.002 ** |
|  |  | Log RMSSD | -143.9605 | 42.6645 | -3.3742 | 0.001 *** |
|  |  | Log SDNN | -257.0395 | 53.8375 | -4.7744 | <0.001 *** |

**Table 4.**
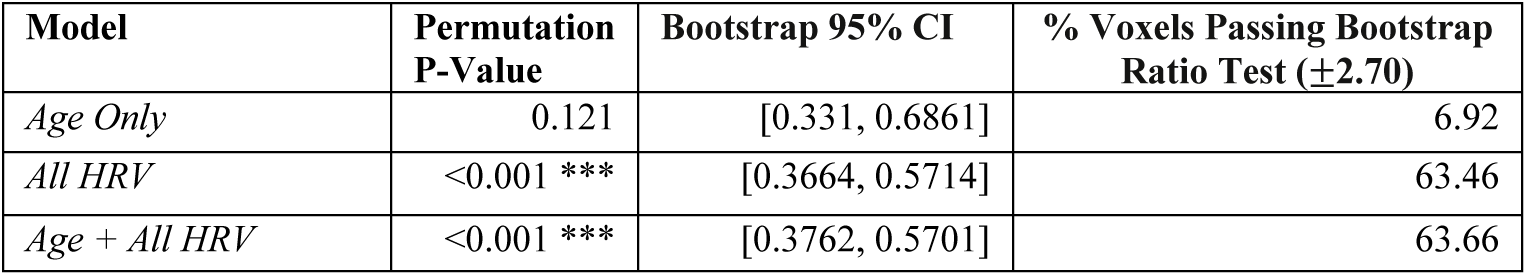
MIDUS Refresher PLS Permutation and Bootstrap Results.

| Model | Permutation P-Value | Bootstrap 95% CI | % Voxels Passing Bootstrap Ratio Test ($\pm 2.70$) |
| --- | --- | --- | --- |
| Age Only | 0.121 | [0.331, 0.6861] | 6.92 |
| All HRV | <0.001 *** | [0.3664, 0.5714] | 63.46 |
| Age + All HRV | <0.001 *** | [0.3762, 0.5701] | 63.66 |

**Table 5.** Within-person analyses from MIDUS Core and MIDUS Refresher (Ref)

| Pred. | Intercept | | Person Centered (PC) vs Mean | $\beta$ | | Std. Error | | T-value | | P-Value | |
| --- | --- | --- | --- | --- | --- | --- | --- | --- | --- | --- | --- |
|  | Core | Ref |  | Core | Ref | Core | Ref | Core | Ref | Core | Ref |
| Log SDNN | 0.1572 | 0.1623 | Person Centered | 0.0087 | 0.0071 | 0.0025 | 0.0018 | 3.49 | 3.95 | <0.001 *** | <0.001 *** |
|  |  |  | Mean | 0.0416 | 0.0511 | 0.0131 | 0.0139 | 3.17 | 3.69 | 0.002 ** | <0.001 *** |
| Log RMSSD | 0.2341 | 0.2480 | Person Centered | 0.0045 | 0.0046 | 0.0025 | 0.0018 | 1.81 | 2.64 | 0.072 . | 0.009 ** |
|  |  |  | Mean | 0.0255 | 0.0338 | 0.0107 | 0.0117 | 2.39 | 2.9 | 0.018 * | 0.005 ** |
| Log LF | 0.1867 | 0.1833 | Person Centered | 0.0091 | 0.0072 | 0.0027 | 0.0015 | 3.42 | 4.68 | <0.001 *** | <0.001 *** |
|  |  |  | Mean | 0.0219 | 0.0245 | 0.0064 | 0.0057 | 3.45 | 4.32 | <0.001 *** | <0.001 *** |
| Log HF | 0.2460 | 0.2645 | Person Centered | 0.0016 | 0.0028 | 0.0024 | 0.0018 | 0.65 | 1.57 | 0.516 | 0.12 |
|  |  |  | Mean | 0.0137 | 0.0163 | 0.0064 | 0.0059 | 2.15 | 2.77 | 0.034 * | 0.007 ** |
| Age | 0.3198 | 0.3560 | NA | -0.0007 | -0.0010 | 0.0010 | 0.0007 | -0.76 | -1.86 | 0.448 | 0.066 . |

### HRV Associations with BOLD Signal Variability

For HRV-only PLS models, 93.9% of voxels showed negative voxel saliences in both Refresher and Core datasets, indicating reliability in HRV-BOLD associations across samples.

#### Core Results

PLS analysis of HRV measures (SDNN, RMSSD, LF, HF) revealed significant associations with SD_BOLD_ (permutation p-value = 0.018; **Tables 1, 2**). The first latent variable explained 97.7% of the covariance structure. Remarkably, 98.3% of brain voxels showed negative voxel weights. Individual HRV predictors showed the following negative associations with brain scores: SDNN (β(114) = -156.74, t = -3.27, p = 0.001), RMSSD (β(114) = -73.80, t = -2.10, p = 0.038), LF (β(114) = -68.99, t = -3.26, p = 0.001), and HF (β(114) = - 38.86, t = -1.97, p = 0.052). The negative beta coefficients, combined with predominantly negative voxel weights, indicate positive associations between all HRV measures and SD_BOLD_. Bootstrap analysis revealed that 31.5% of voxels passed significance testing. Greater than 99% of voxels passing bootstrap ratio tests exhibited negative voxel weights.

#### Refresher Results

PLS analysis of HRV measures once again proved to be highly significant, with permutation p-value < 0.001 (**Tables 3, 4**). The first latent variable explained 97.8% of covariance, and individual HRV predictors demonstrated strong negative associations with brain scores: SDNN (β(101) = -257.20, t = -4.76, p < 0.001), RMSSD (β(101) = -143.96, t = -3.36, p = 0.001), LF (β(101) = -123.88, t = -5.85, p < 0.001), and HF (β(101) = -68.76, t = -3.24, p = 0.002). Negative associations between HRV and brain scores, paired with majority negative voxel weights suggests that across the brain, HRV is associated with increases in SD_BOLD_. 96.3% of voxels showed negative weights, with 63.5% passing bootstrap significance testing. Greater than 99% of voxels which passed bootstrap ratio tests also showed negative voxel weights.

### Joint Age and HRV Models

#### Core Results

When age and HRV measures were modeled simultaneously, joint analyses remained significant (permutation p-value = 0.019; **Tables 1, 2**; **Figs. 3, 4**). The first latent variable explained 89.7% of covariance, with 98% negative voxel weights. Critically, individual HRV predictors demonstrated strong negative associations with brain scores from the first latent variable (**Fig. 3**): SDNN (β(114) = -156.64, t = -3.26, p = 0.002), RMSSD (β(114) = -73.41, t = 2.03, p = 0.045), LF (β(114) = -69.14, t = -3.25, p = 0.002), and HF (β(114) = -38.71, t = -1.95, p = 0.054), while age was not significantly associated with brain scores from the first latent variable (β(114) = 2.19, t = 0.68, p = 0.498). The second latent variable explained 8.7% of covariance, with only 53% negative voxel weights. Age was strongly associated with brain scores from the second latent variable (β(114) = 5.07, t = 7.70, p < 0.001), while none of the HRV measures were associated with brain scores across the second latent variable: SDNN (β(114) = 5.10, t = 0.40, p = 0.688), RMSSD (β(114) = 12.18, t = 1.35, p = 0.179), LF (β(114) = -3.00, t = -0.54, p = 0.593), and HF (β = 5.07, t = 1.00, p = 0.319). We found that 31.5% of voxels passed bootstrap ratio significance testing, and greater than 99% of those voxels passing bootstrap ratio tests exhibited negative voxel weights.

**Figure 3:**
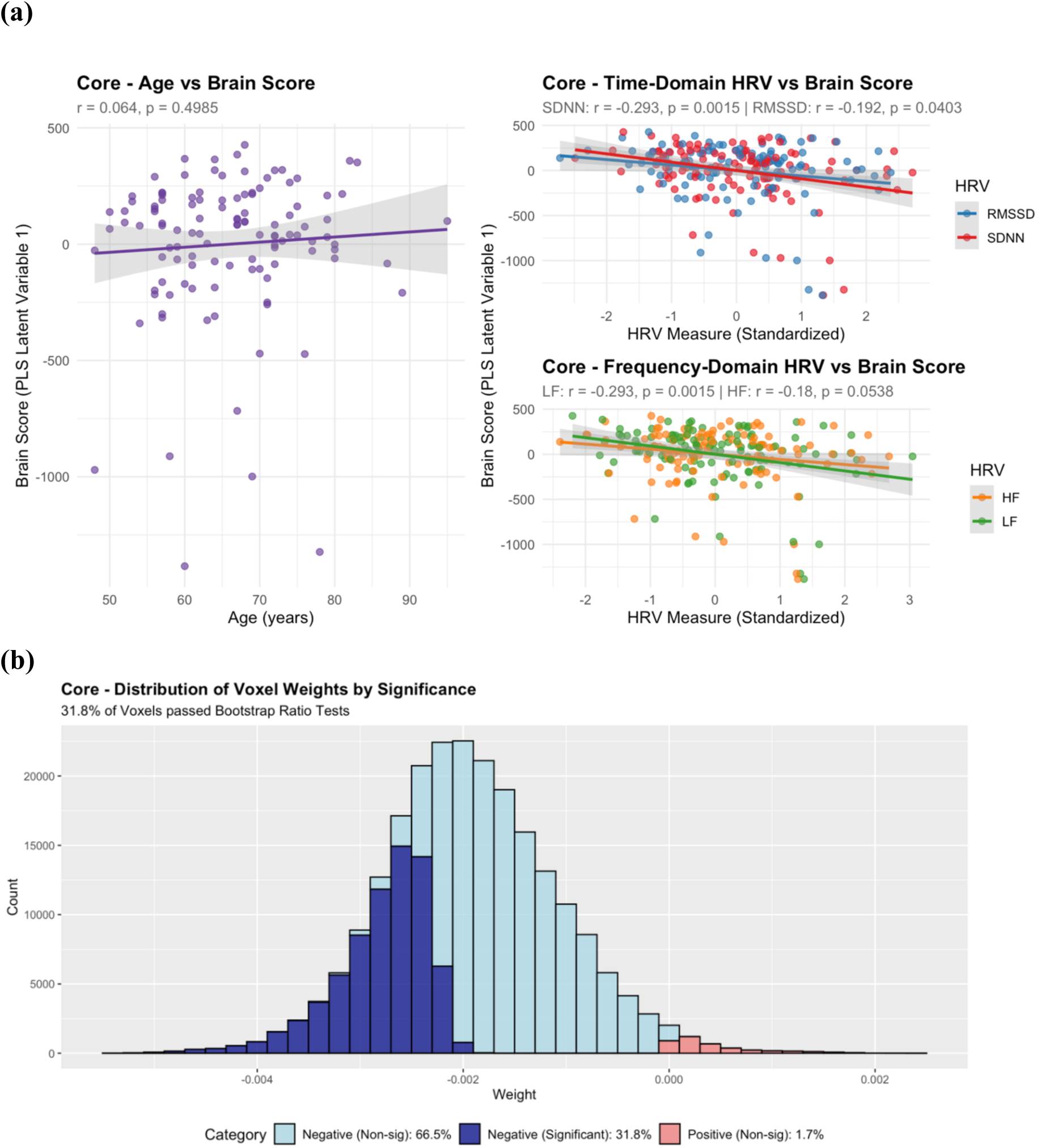
**(a) MIDUS Core Latent Variable 1 Brain Scores vs Behavioral Variables** displaying regression relationships between age, HRV, and brain scores of joint PLS Age and HRV models. Latent variable 1 captures the dominant pattern of covariation between SD_BOLD_ and predictor variables, explaining 89.7% of the total covariance and predominantly reflecting HRV-related associations with SD_BOLD_. Higher brain scores indicate that a subject more closely expresses the multivariate pattern displayed by voxel weights in (b). **(b) MIDUS Core Histogram of Voxel Saliences (Latent Variable 1)** from joint PLS models. Voxel saliences are colored by their relative voxel weight (negative vs positive) and whether they passed bootstrap ratio correction at ± 2.7. Predominantly negative voxel weights suggests that subjects with higher brain scores show lower SD_BOLD_.

**Figure 4:**
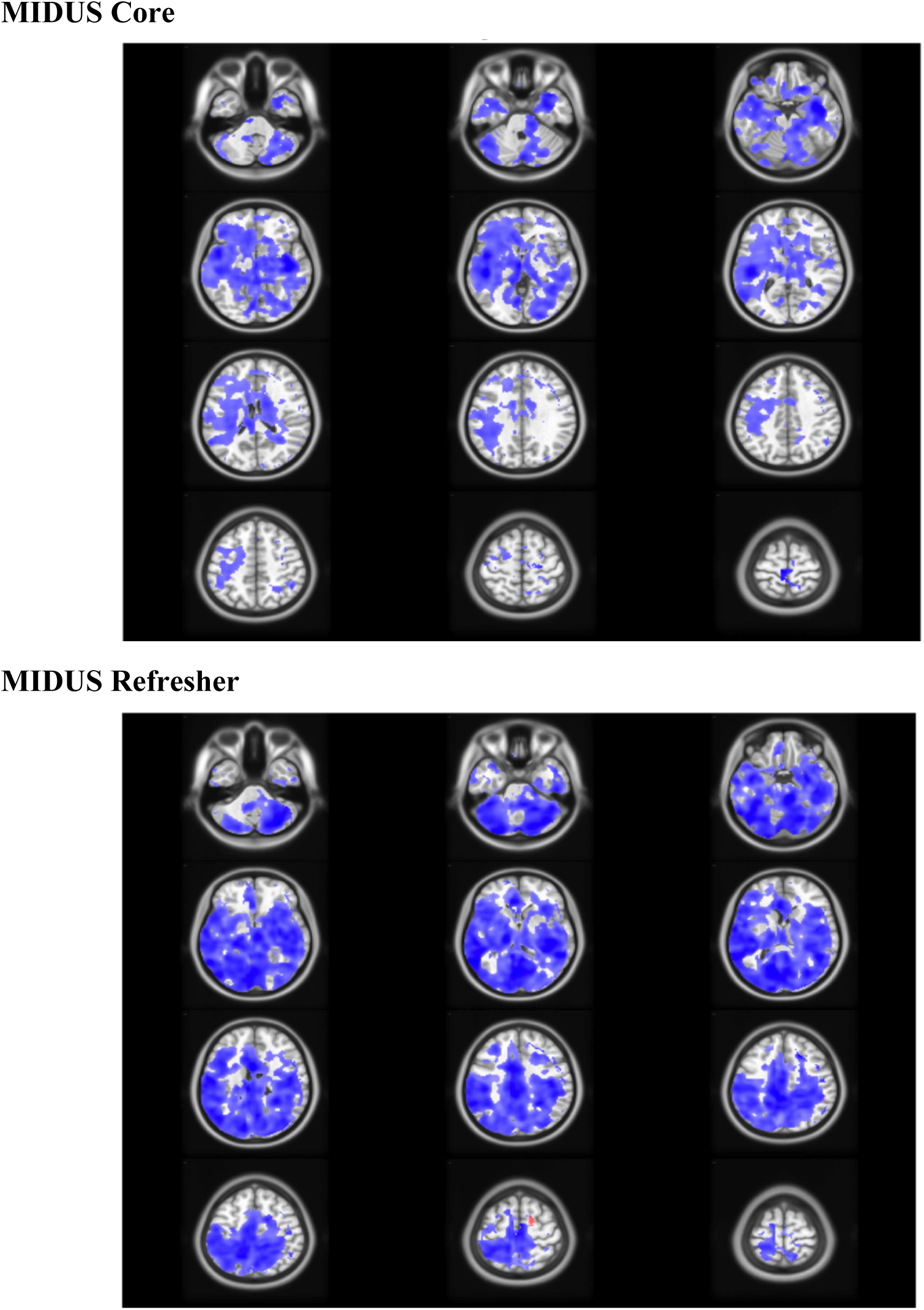
**Brain Salience Weights (Latent Variable 1)** of voxels passing *Bootstrap Ratio Tests* for HRV and Age joint PLS models in Core and Refresher Samples. Latent variable 1 captures the dominant pattern of covariation between SD_BOLD_ and the predictor variables, explaining 89.7% (Core) and 94.2% (Refresher) of the total covariance and predominantly reflecting HRV-related associations with SD_BOLD_. Colored regions display voxel weights which passed bootstrap ratio correction at ± 2.7, colored by voxel saliences (blue = negative, red = positive). Blue regions indicate positive associations between HRV and SD_BOLD_, whereas red regions indicate negative associations.

#### Refresher Results

Similarly, in MIDUS Refresher, joint models remained significant (permutation p-value < 0.001; **Tables 3, 4**; **Figs. 4, 5**). The first latent variable explained 94.2% of covariance, with 96% negative voxel weights. Individual HRV predictors demonstrated strong negative associations with brain scores from the first latent variable (**Fig. 5**): SDNN (β(101) = - 257.04, t = -4.77, p < 0.001), RMSSD (β(101) = -143.96, t = -3.37, p = 0.001), LF (β(101) = - 123.73, t = -5.86, p < 0.001), and HF (β(101) = -68.83, t = -3.26, p = 0.002), while age was not significantly associated with brain scores from the first latent variable (β(101) = 3.18, t = 1.27, p = 0.207). Alternatively, the second latent variable explained 3.8% of the covariance, with only 42% negative voxel weights. Age was strongly associated with brain scores from the second latent variable (β(101) = 4, t = 5.65, p < 0.001), while none of the HRV measures were associated with brain scores across the second latent variable: SDNN (β(101) = 9.25, t = 0.49, p = 0.630), RMSSD (β(101) = -2.51, t = -0.17, p = 0.863), LF (β(101) = 9.00, t = 1.15, p = 0.252), and HF (β(101) = -2.58, t = -0.36, p = 0.72). Bootstrap analysis revealed that 63% of voxels passed significance testing, with greater than 99% passing bootstrap ratio tests exhibited negative voxel weights.

**Figure 5:**
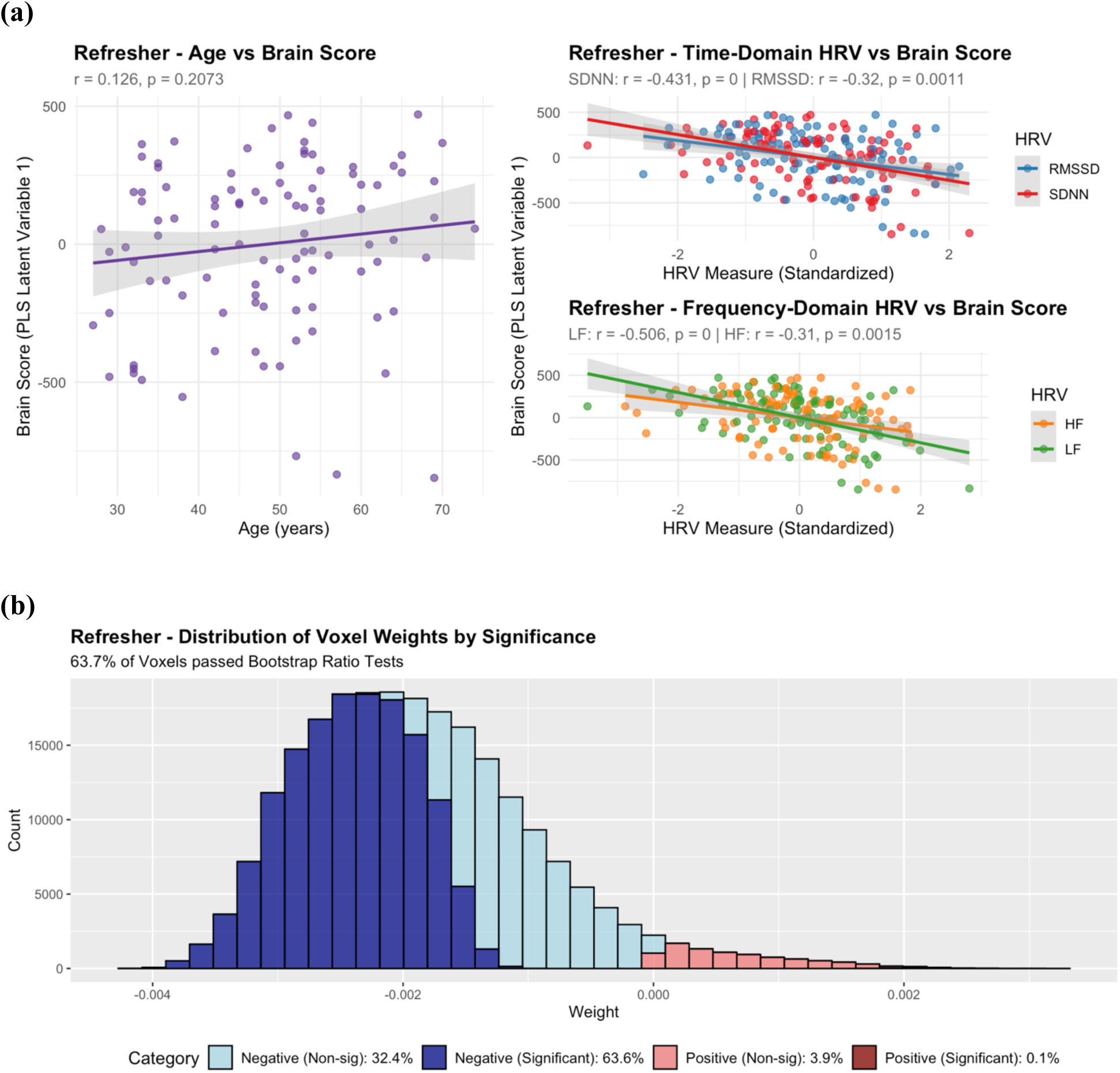
(a) MIDUS Refresher Latent Variable 1 Brain Scores vs Age and HRV displaying regression relationship between age, HRV, and brain scores of joint PLS Age and HRV models. Latent variable 1 captures the dominant pattern of covariation between SD_BOLD_ and predictor variables, explaining 94.2% of the total covariance and predominantly reflecting HRV-related associations with SD_BOLD_. Higher brain scores indicate that a subject more closely expresses the multivariate pattern displayed by voxel weights in (b). **(b) MIDUS Refresher Histogram of Voxel Saliences (Latent Variable 1)** from joint PLS models. Voxel saliences are colored by their relative voxel weight (negative vs positive) and whether they passed bootstrap ratio correction at ± 2.7. Predominantly negative voxel weights suggests that subjects with higher brain scores show lower SD_BOLD_.

Results from both samples suggest that age explains dramatically less variance in SD_BOLD_ than HRV across measures. When HRV is included in joint models, the directional negative effects of age on SD_BOLD_ disappear, as shown in the difference in voxel weight proportions between latent variables from the age-only model and the second latent variable from the joint models.

### Spatial Concordance Analysis

Concordance analyses examining the overlap between age-related and HRV-related spatial patterns in the first latent variable revealed consistent findings across datasets. In MIDUS Core, 69.1% of brain voxels showed concordant negative effects (regions with age-related decreases in SD_BOLD_ also showing HRV-related increases), while 0.8% showed concordant positive effects, and 30.1% showed opposite directional effects. MIDUS Refresher showed very similar concordance, with 67.0% of voxels demonstrating concordant negative effects, 4.0% showing concordant positive effects, and 29.0% showing opposite patterns. These concordance patterns support the interpretation that brain regions showing age-related reductions in SD_BOLD_ are predominantly the same regions showing HRV-related increases in variability, consistent with the hypothesis that cardiovascular factors may be partially responsible for apparent age-related changes in neural signal dynamics.

### Within-person Sliding Window Analyses

#### Core Results

Across all measures, we found strong positive correlations between mean HRV and SD_BOLD_ (**Fig. 6**): SDNN (β(114) = 0.042, t = 3.17, p = 0.002), RMSSD (β(114) = 0.026, t = 2.39, p = 0.018), HF (β(114) = 0.014, t = 2.15, p = 0.034) and LF (β(114) = 0.022, t = 3.45, p < 0.001) HRV. We found non-significant associations between age and average SD_BOLD_ (β(114) = -0.0007, t = -0.76, p = 0.448). We also found significant positive correlations between SD_BOLD_ and person-centered SDNN (β(114) = 0.009, t = 3.49, p < 0.001) and LF (β(114) = 0.009, t = 3.42, p < 0.001), but not HF (β(114) = 0.002, t = 0.65, p = 0.516) HRV nor RMSSD (β(114) = 0.005, t = 1.81, p = 0.072).

**Figure 6:**
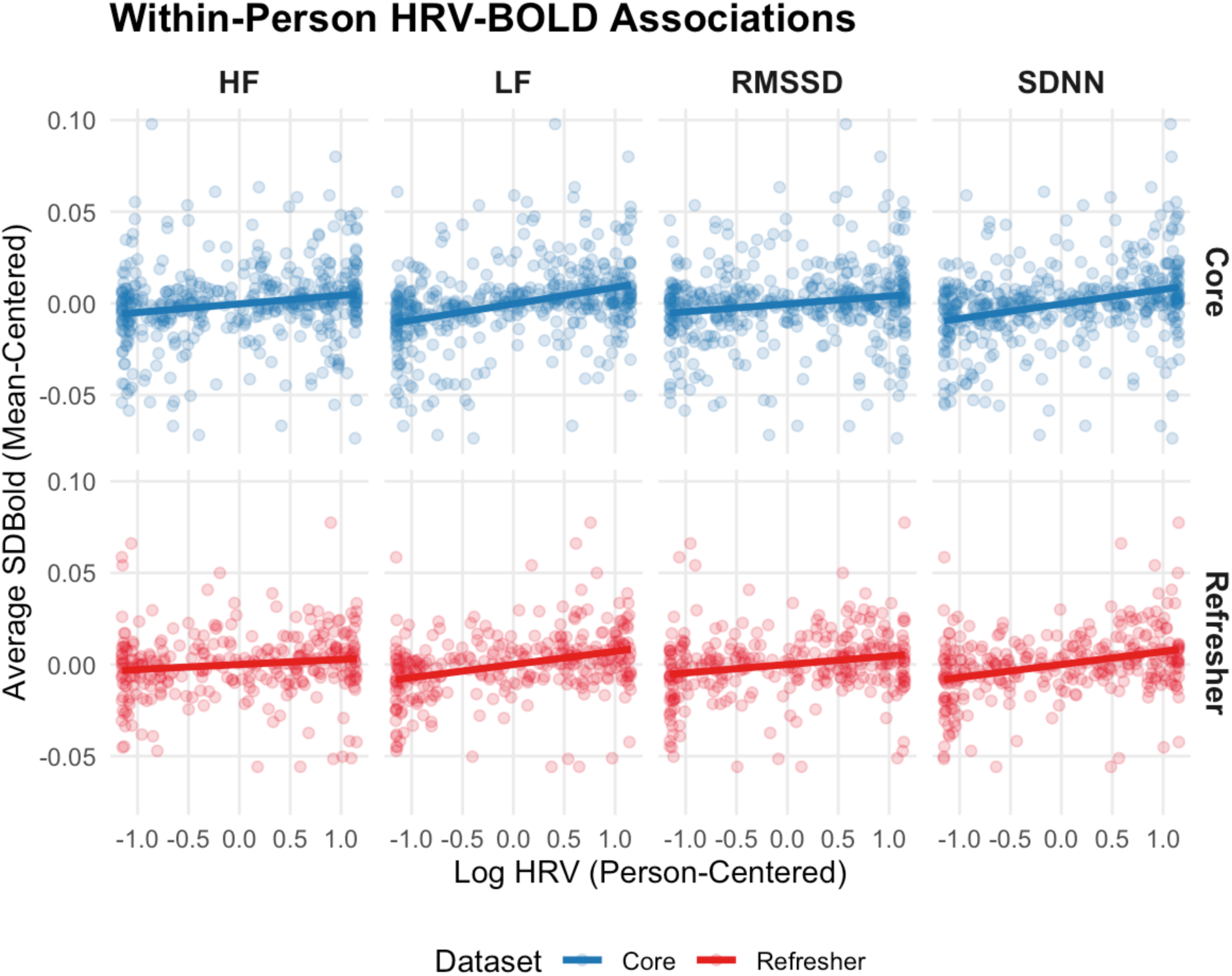
Within-Person Associations Between SD_BOLD_ and HRV. We calculated HRV and SD_BOLD_ within three sliding windows for each subject and then centered each window based on subject mean and standard deviation. Higher Person-Centered Log HRV indicates that a given window shows higher HRV when compared to the subject’s average HRV. Higher average SD_BOLD_ (mean-centered) indicates that a given window has higher SD_BOLD_ when compared to a given subject’s average. Positive associations suggest that within participants, windows with higher HRV have higher SD_BOLD._

#### Refresher Results

Similarly, in the refresher sample we observed significant positive correlations between mean HRV and SD_BOLD_ (**Fig. 6**): SDNN (β(101) = 0.051, t = 3.69, p < 0.001), RMSSD (β(101) = 0.034, t = 2.9, p = 0.005), HF (β(101) = 0.014, t = 2.77, p = 0.007) and LF (β(101) = 0.025, t = 4.32, p < 0.001) HRV. We found marginally significant negative associations between age and average SD_BOLD_ (β(101) = -0.001, t = -1.86, p = 0.066). We also found significant positive correlations between SD_BOLD_ and person-centered SDNN (β(101) = 0.009, t = 3.95, p < 0.001), RMSSD (β(101) = 0.005, t = 2.64, p = 0.009), and LF (β(101) = 0.007, t = 4.68, p < 0.001), but not HF (β(101) = 0.003, t = 1.57, p = 0.12) HRV.

Across all measures, mean HRV was positively associated with SD_BOLD_, validating the results observed through PLS modeling. These robust within-person associations suggest that across subjects, windows with higher variance simultaneously had higher SD_BOLD_, regardless of subject-specific baseline levels of HRV or SD_BOLD_.

## Discussion

The present study examined associations between chronological age, different measures of heart rate variability (HRV) and BOLD signal variability (SD_BOLD_) during resting-state fMRI across two independent datasets, providing novel insights into cardiovascular contributions to neural signal dynamics. Within the MIDUS Core sample at MIDUS 3, age was not associated with SDNN, LF nor HF HRV, and age was positively associated with RMSSD (though this association did not pass false discovery rate (FDR) correction). Within the MIDUS Refresher sample at the MIDUS Refresher 1 timepoint, SDNN, LF, and HF HRV were negatively associated with age after FDR correction, but age and RMSSD were not significantly associated. Across both Core and Refresher datasets, we observed robust positive associations between multiple HRV indices and SD_BOLD_, with effect sizes consistently exceeding those for chronological age alone. Through PLS analyses in both Core and Refresher samples, permutation tests revealed significant associations between HRV and SD_BOLD_ (Core: p = 0.018, Refresher: p < 0.001) despite age-only models being insignificant (Core: p = 0.201, Refresher: p = 0.121). Spatial concordance analyses revealed that brain regions showing age-related decreases in SD_BOLD_ were predominantly the same voxels showing HRV-related increases in variability. In the Refresher sample, 67% of voxels demonstrated this inverse relationship, and 69% in the Core sample. Joint modeling approaches consistently demonstrated that HRV measures explained larger proportions of variance in BOLD signal patterns than chronological age across both datasets. Within-person sliding window analyses revealed strong positive correlations between within-person mean centered HRV and SD_BOLD_. Findings suggest that windows with higher HRV also exhibited higher SD_BOLD_, regardless of participant baseline levels in SD_BOLD_ or HRV.

Our findings reinforce the complexity of age-HRV relationships across the life course. Negative associations between age and HRV within MIDUS Refresher echo those of prior literature which suggests that HRV is negatively associated with age (Almeida-Santos et al., 2016; Antelmi et al., 2004), with decreases plateauing around age 50. Our findings from MIDUS Core showed no significant associations between SDNN, LF, and HF HRV with age. Because MIDUS Core (mean age=65.9 years, SD=9.1) comprises a significantly older sample than MIDUS Refresher, positive associations between RMSSD and age in the Core sample are somewhat consistent with past literature. Research suggests that RMSSD may increase after age 50 (Almeida-Santos et al., 2016), and that the majority of age-related declines in HRV are seen in younger adulthood (Antelmi et al., 2004). MIDUS Core comprises a significantly older cohort than MIDUS Refresher (MR Age: M=48.5, SD=11.6, range=26-76; M3 Age: M=65.9, SD=9.1, range=48-95) which may partially explain these differences in significance across samples. Additionally, associations between RMSSD and age were non-significant after FDR correction in both Core and Refresher samples, suggesting that these effects may be less robust than the consistent age-related declines observed in SDNN, LF, and HF HRV within the Refresher sample.

Our research provides the first direct PLS-based evidence linking HRV to the SD_BOLD_ measure introduced by Garrett et al. (2010), extending previous cardiovascular confound research beyond the RSFA methodology to demonstrate cardiovascular influences across a broader frequency range of neural signal dynamics. These findings align with Tsvetanov et al. (2021), who argued that age related decreases in RSFA are explained by cardiovascular and cerebrovascular factors. Our study extends this work by directly examining the SD_BOLD_ measure introduced by Garrett et al. (2010), rather than the RSFA measure that has been the focus of previous studies. The robust HRV-BOLD variability relationships we observe using the SD_BOLD_ methodology suggest that correlations between HRV and BOLD variability extend beyond the low-frequency oscillations (0.01-0.08 Hz) emphasized by RSFA. These correlations may reflect either the influence of vascular effects on SD_BOLD,_ or the influence of neural factors (like arousal) or another third variable on both HRV and BOLD variability. Importantly, our joint modeling approach revealed that when both age and HRV were included simultaneously, HRV consistently explained larger proportions of variance in SD_BOLD_ patterns. This finding suggests that chronological age per se may be less important in explaining differences in SD_BOLD_ than the HRV changes that typically accompany aging. Our observed within-person associations between HRV and SD_BOLD_ provide further strong support for this.

The present findings provide a nuanced perspective on the debate regarding vascular confounds in BOLD variability measures (Garrett et al., 2017; Tsvetanov et al., 2021). The BOLD signal is a convolution of both neural and vascular components, potentially leading to cardiovascular components being picked up in SD_BOLD_ even after extensive preprocessing. Garrett et al. (2017) argued that age differences in SD_BOLD_ were maintained after adjusting for multiple vascular factors. Garrett et al. measured vascular parameters such as baseline cerebral blood flow and BOLD cerebrovascular reactivity, which were obtained during a dedicated hypercapnia session that took place shortly after resting fixation runs used to estimate SD_BOLD_. After collecting resting fixation runs, participants were allowed to get out of the fMRI scanner for a break, then put back into the scanner lying down for the hypercapnia session used to estimate vascular parameters. Notably, their vascular control analyses did not include simultaneous HRV measures, which our study demonstrates to be particularly potent predictors of SD_BOLD_. Our HRV measures were acquired simultaneously with the resting-state fMRI data, ensuring temporal alignment between cardiovascular and BOLD signal dynamics. This simultaneous acquisition may provide a more direct assessment of the cardiovascular contributions to SD_BOLD_ than vascular measures obtained in separate scanning sessions.

Our results suggest that the specific cardiovascular measure examined matters significantly. While Garrett et al. (2017) controlled for measures including cerebral blood flow and BOLD cerebrovascular reactivity acquired during separate sessions, HRV may capture unique aspects of autonomic nervous system function that are more directly relevant to SD_BOLD_. HRV reflects the complex interplay between sympathetic and parasympathetic nervous system activity, which directly influences cerebrovascular dynamics and neurovascular coupling mechanisms that underlie the BOLD response (Zhang, 2007). Since the Refresher sample showed strong negative correlations between age and HRV, while the Core sample did not, our findings suggest that when conducting between-subjects tests with SD_BOLD_, controlling for HRV may be most important in younger samples (25-50), where HRV is often in active decline.

It is important to consider that previous literature analyzing age and SD_BOLD_ has used age as a categorical variable, with significant gaps (∼20 years) between younger vs. older participants (Garrett et al., 2010, 2013, 2017). Both samples examined in this project comprised a large continuous age range (Core: ages 26-76; Refresher: ages 48-95); hence we opted to use age as a continuous variable. Treating age as a continuous variable allowed for more granular assessment of age-related changes in SD_BOLD_. However, future research should address these findings with non-linear alternatives to PLS (e.g., kernel PLS) to determine whether SD_BOLD_ may decline non-linearly with increases in continuous age. Additionally, the Core sample comprised a much older population than previous literature finding that SD_BOLD_ declines with age, potentially explaining the lack of associations between age and SD_BOLD_ seen in the Core.

Several limitations and future directions should be considered. While simultaneous acquisition of HRV and fMRI data represents a methodological strength, photoplethysmography may be less precise than electrocardiography for estimating beat-to-beat intervals. Additionally, non-negative associations between age and HRV in MIDUS Core may partly be due to selection effects. The older individuals in the longitudinal MIDUS Core cohort are in their 80s and 90s and can travel to participate in MIDUS fMRI scans, lie flat on their back for an hour, and in general may be healthier comparatively to others their age, reflecting the observed higher RMSSD among these older participants. Participants with poor cardiovascular function (i.e., low HRV) may also have passed away, leading to survivorship bias and non-random attrition in our sample. Lastly, non-significant age-SD_BOLD_ associations via PLS permutation tests poses a limitation, constraining our ability to conclusively demonstrate that controlling for HRV eliminates age effects. This limitation necessitates replication in samples with stronger baseline age-SD_BOLD_ relationships.

Collectively, we find BOLD signal variability is robustly associated with heart rate variability, with effect sizes that consistently exceed those for chronological age. Our results strongly indicate that HRV should be measured and controlled for when analyzing BOLD signal variability.

## Acknowledgements

We are extremely grateful to the Midlife in the U.S. (MIDUS) participants and staff for their contributions to the study without which this research would not be possible. Publicly available data from the MIDUS study was used for this research. Since 1995, the MIDUS study has been funded by the following: the John D. and Catherine T. MacArthur Foundation Research Network, and the National Institute on Aging (P01-AG020166, U19-AG051426, U01-AG077928). Data collection for the MIDUS Neuroscience Project received partial support through a core grant awarded to the Waisman Center at the University of Wisconsin-Madison by the National Institute of Child Health and Human Development (P50HD105353). The authors also acknowledge support from the Fulbright-Mitacs Globalink Research Program, which supported analysis and collaboration conducted at McGill University in Canada. Additional support was provided by a grant from the Natural Sciences and Engineering Research Council of Canada (NSERC) Discovery Research program (RGPIN-2022-05399) and Supplement (DGECR-2022-00321) to BSK, and computing resources from Calcul Quebec and the Digital Research Alliance of Canada.

## Statements and Declarations

### Ethics Approval

All procedures were approved by the University of Wisconsin-Madison Institutional Review Board. MIDUS IRB Protocol Number – Neuroscience (MRI): 2016-0054.

### Data Availability

Raw MIDUS neuroimaging and psychophysiological data publicly available through restricted access procedures. Instructions for requesting data and obtaining access can be found here: https://midus.wisc.edu/?page_id=98.

### Informed Consent

Informed consent was obtained from all participants included in the study.

### Availability of Data and Material

Raw MIDUS neuroimaging and psychophysiological data are publicly available through restricted access procedures. Instructions for requesting data and obtaining access can be found here: https://midus.wisc.edu/?page_id=98.

## References

Abraham, Alexandre, Fabian Pedregosa, Michael Eickenberg, Philippe Gervais, Andreas Mueller, Jean Kossaifi, Alexandre Gramfort, Bertrand Thirion, and Gael Varoquaux. 2014. “Machine Learning for Neuroimaging with Scikit-Learn.” Frontiers in Neuroinformatics 8. 10.3389/fninf.2014.00014.

Almeida-Santos, M. A., Barreto-Filho, J. A., Oliveira, J. L. M., Reis, F. P., da Cunha Oliveira, C. C., & Sousa, A. C. S. (2016). Aging, heart rate variability and patterns of autonomic regulation of the heart. Archives of Gerontology and Geriatrics, 63, 1–8. 10.1016/j.archger.2015.11.011

Antelmi, I., De Paula, R. S., Shinzato, A. R., Peres, C. A., Mansur, A. J., & Grupi, C. J. (2004). Influence of age, gender, body mass index, and functional capacity on heart rate variability in a cohort of subjects without heart disease. American Journal of Cardiology, 93(3), 381–385. 10.1016/j.amjcard.2003.09.065

Avants, B. B., C. L. Epstein, M. Grossman, and J. C. Gee. 2008. “Symmetric Diffeomorphic Image Registration with Cross-Correlation: Evaluating Automated Labeling of Elderly and Neurodegenerative Brain.” Medical Image Analysis 12 (1): 26–41. 10.1016/j.media.2007.06.004.

Behzadi, Yashar, Khaled Restom, Joy Liau, and Thomas T. Liu. 2007. “A Component Based Noise Correction Method (CompCor) for BOLD and Perfusion Based fMRI.” NeuroImage 37 (1): 90–101. 10.1016/j.neuroimage.2007.04.042.

Cox, Robert W., and James S. Hyde. 1997. “Software Tools for Analysis and Visualization of fMRI Data.” NMR in Biomedicine 10 (4-5): 171–78. 10.1002/(SICI)1099-1492(199706/08)10:4/5<171::AID-NBM453>3.0.CO;2-L.

Esteban, O., Markiewicz, C. J., Blair, R. W., Moodie, C. A., Isik, A. I., Erramuzpe, A., Kent, J. D., Goncalves, M., DuPre, E., Snyder, M., Oya, H., Ghosh, S. S., Wright, J., Durnez, J., Poldrack, R. A., & Gorgolewski, K. J. (2019). fMRIPrep: A robust preprocessing pipeline for functional MRI. Nature Methods, 16(1), 111–116. 10.1038/s41592-018-0235-4

Esteban, Oscar, Christopher Markiewicz, Ross W Blair, Craig Moodie, Ayse Ilkay Isik, Asier Erramuzpe Aliaga, James Kent, et al. 2018. “fMRIPrep: A Robust Preprocessing Pipeline for Functional MRI.” Nature Methods. 10.1038/s41592-018-0235-4.

Esteban, Oscar, Ross Blair, Christopher J. Markiewicz, Shoshana L. Berleant, Craig Moodie, Feilong Ma, Ayse Ilkay Isik, et al. 2018. “fMRIPrep 23.0.0.” Software. 10.5281/zenodo.852659.

Evans, AC, AL Janke, DL Collins, and S Baillet. 2012. “Brain Templates and Atlases.” NeuroImage 62 (2): 911–22. 10.1016/j.neuroimage.2012.01.024.

Fonov, VS, AC Evans, RC McKinstry, CR Almli, and DL Collins. 2009. “Unbiased Nonlinear Average Age-Appropriate Brain Templates from Birth to Adulthood.” NeuroImage 47, Supplement 1: S102. 10.1016/S1053-8119(09)70884-5.

Garrett, D. D., Kovacevic, N., McIntosh, A. R., & Grady, C. L. (2010). Blood Oxygen Level-Dependent Signal Variability Is More than Just Noise. The Journal of Neuroscience, 30(14), 4914. 10.1523/JNEUROSCI.5166-09.2010

Garrett, D. D., Kovacevic, N., McIntosh, A. R., & Grady, C. L. (2013). The Modulation of BOLD Variability between Cognitive States Varies by Age and Processing Speed. Cerebral Cortex, 23(3), 684–693. 10.1093/cercor/bhs055

Garrett, D. D., Lindenberger, U., Hoge, R. D., & Gauthier, C. J. (2017). Age differences in brain signal variability are robust to multiple vascular controls. Scientific Reports, 7(1), 10149. 10.1038/s41598-017-09752-7

Gorgolewski, K., C. D. Burns, C. Madison, D. Clark, Y. O. Halchenko, M. L. Waskom, and S. Ghosh. 2011. “Nipype: A Flexible, Lightweight and Extensible Neuroimaging Data Processing Framework in Python.” Frontiers in Neuroinformatics 5: 13. 10.3389/fninf.2011.00013.

Gorgolewski, Krzysztof J., Oscar Esteban, Christopher J. Markiewicz, Erik Ziegler, David Gage Ellis, Michael Philipp Notter, Dorota Jarecka, et al. 2018. “Nipype.” Software. 10.5281/zenodo.596855.

Grady, C. L., & Garrett, D. D. (2018). Brain signal variability is modulated as a function of internal and external demand in younger and older adults. NeuroImage, 169, 510–523. 10.1016/j.neuroimage.2017.12.031

Greve, Douglas N, and Bruce Fischl. 2009. “Accurate and Robust Brain Image Alignment Using Boundary-Based Registration.” NeuroImage 48 (1): 63–72. 10.1016/j.neuroimage.2009.06.060.

Guitart-Masip, M., Salami, A., Garrett, D., Rieckmann, A., Lindenberger, U., & Bäckman, L. (2016). BOLD Variability is Related to Dopaminergic Neurotransmission and Cognitive Aging. Cerebral Cortex, 26(5), 2074–2083. 10.1093/cercor/bhv029

Jenkinson, Mark, and Stephen Smith. 2001. “A Global Optimisation Method for Robust Affine Registration of Brain Images.” Medical Image Analysis 5 (2): 143–56. 10.1016/S1361-8415(01)00036-6.

Jenkinson, Mark, Peter Bannister, Michael Brady, and Stephen Smith. 2002. “Improved Optimization for the Robust and Accurate Linear Registration and Motion Correction of Brain Images.” NeuroImage 17 (2): 825–41. 10.1006/nimg.2002.1132.

Kannurpatti, S. S., & Biswal, B. B. (2008). Detection and scaling of task-induced fMRI-BOLD response using resting state fluctuations. NeuroImage, 40(4), 1567–1574. 10.1016/j.neuroimage.2007.09.040

Krishnan, A., Williams, L. J., McIntosh, A. R., & Abdi, H. (2011). Partial Least Squares (PLS) methods for neuroimaging: A tutorial and review. Multivariate Decoding and Brain Reading, 56(2), 455–475. 10.1016/j.neuroimage.2010.07.034

Lanczos, C. 1964. “Evaluation of Noisy Data.” Journal of the Society for Industrial and Applied Mathematics Series B Numerical Analysis 1 (1): 76–85. 10.1137/0701007.

Logothetis, N. What we can do and what we cannot do with fMRI. Nature 453, 869–878 (2008). 10.1038/nature06976

Makowski, D., Pham, T., Lau, Z. J., Brammer, J. C., Lespinasse, F., Pham, H., Schölzel, C., & Chen, S. A. (2021). NeuroKit2: A Python toolbox for neurophysiological signal processing. Behavior Research Methods, 53(4), 1689–1696. 10.3758/s13428-020-01516-y

McIntosh, A. R., Kovacevic, N., & Itier, R. J. (2008). Increased Brain Signal Variability Accompanies Lower Behavioral Variability in Development. PLOS Computational Biology, 4(7), e1000106. 10.1371/journal.pcbi.1000106

Millar, P. R., Petersen, S. E., Ances, B. M., Gordon, B. A., Benzinger, T. L. S., Morris, J. C., & Balota, D. A. (2020). Evaluating the Sensitivity of Resting-State BOLD Variability to Age and Cognition after Controlling for Motion and Cardiovascular Influences: A Network-Based Approach. Cerebral Cortex, 30(11), 5686–5701. 10.1093/cercor/bhaa138

Patriat, Rémi, Richard C. Reynolds, and Rasmus M. Birn. 2017. “An Improved Model of Motion-Related Signal Changes in fMRI.” NeuroImage 144, Part A (January): 74–82. 10.1016/j.neuroimage.2016.08.051.

Power, Jonathan D., Anish Mitra, Timothy O. Laumann, Abraham Z. Snyder, Bradley L. Schlaggar, and Steven E. Petersen. 2014. “Methods to Detect, Characterize, and Remove Motion Artifact in Resting State fMRI.” NeuroImage 84 (Supplement C): 320–41. 10.1016/j.neuroimage.2013.08.048.

Pruim, Raimon H. R., Maarten Mennes, Daan van Rooij, Alberto Llera, Jan K. Buitelaar, and Christian F. Beckmann. 2015. “ICA-AROMA: A Robust ICA-Based Strategy for Removing Motion Artifacts from fMRI Data.” NeuroImage 112 (Supplement C): 267–77. 10.1016/j.neuroimage.2015.02.064.

Rieck, J. R., Rodrigue, K. M., Boylan, M. A., & Kennedy, K. M. (2017). Age-related reduction of BOLD modulation to cognitive difficulty predicts poorer task accuracy and poorer fluid reasoning ability. NeuroImage, 147, 262–271. 10.1016/j.neuroimage.2016.12.022

Satterthwaite, Theodore D., Mark A. Elliott, Raphael T. Gerraty, Kosha Ruparel, James Loughead, Monica E. Calkins, Simon B. Eickhoff, et al. 2013. “An improved framework for confound regression and filtering for control of motion artifact in the preprocessing of resting-state functional connectivity data.” NeuroImage 64 (1): 240–56. 10.1016/j.neuroimage.2012.08.052.

Steinberg, S. N., & King, T. Z. (2024). Within-Individual BOLD Signal Variability and its Implications for Task-Based Cognition: A Systematic Review. Neuropsychology review, 34(4), 1115–1164. 10.1007/s11065-023-09619-x

Tegegne, B. S., Man, T., van Roon, A. M., Riese, H., & Snieder, H. (2018). Determinants of heart rate variability in the general population: The Lifelines Cohort Study. Heart Rhythm, 15(10), 1552–1558. 10.1016/j.hrthm.2018.05.006

Tsvetanov, K. A., Henson, R. N. A., & Rowe, J. B. (2020). Separating vascular and neuronal effects of age on fMRI BOLD signals. Philosophical Transactions of the Royal Society B: Biological Sciences, 376(1815), 20190631. 10.1098/rstb.2019.0631

Tsvetanov, K. A., Henson, R. N. A., Jones, P. S., Mutsaerts, H., Fuhrmann, D., Tyler, L. K., Cam-CAN, & Rowe, J. B. (2021). The effects of age on resting-state BOLD signal variability is explained by cardiovascular and cerebrovascular factors. Psychophysiology, 58(7), e13714. 10.1111/psyp.13714

Tsvetanov, K. A., Henson, R. N. A., Tyler, L. K., Davis, S. W., Shafto, M. A., Taylor, J. R., Williams, N., Cam-CAN, & Rowe, J. B. (2015). The effect of ageing on fMRI: Correction for the confounding effects of vascular reactivity evaluated by joint fMRI and MEG in 335 adults. Human Brain Mapping, 36(6), 2248–2269. 10.1002/hbm.22768

Tustison, N. J., B. B. Avants, P. A. Cook, Y. Zheng, A. Egan, P. A. Yushkevich, and J. C. Gee. 2010. “N4itk: Improved N3 Bias Correction.” IEEE Transactions on Medical Imaging 29 (6): 1310–20. 10.1109/TMI.2010.2046908.

Voss, A., Schroeder, R., Heitmann, A., Peters, A., & Perz, S. (2015). Short-Term Heart Rate Variability—Influence of Gender and Age in Healthy Subjects. PLOS ONE, 10(3), e0118308. 10.1371/journal.pone.0118308

Xifra-Porxas, A., Kassinopoulos, M., & Mitsis, G. D. (2021). Physiological and motion signatures in static and time-varying functional connectivity and their subject identifiability. eLife, 10, e62324. 10.7554/eLife.62324

Zang, Y. F., He, Y., Zhu, C. Z., Cao, Q. J., Sui, M. Q., Liang, M., Tian, L. X., Jiang, T. Z., & Wang, Y. F. (2007). Altered baseline brain activity in children with ADHD revealed by resting-state functional MRI. Brain & development, 29(2), 83–91. 10.1016/j.braindev.2006.07.002

Zhang, J. (2007). Effect of Age and Sex on Heart Rate Variability in Healthy Subjects. Journal of Manipulative & Physiological Therapeutics, 30(5), 374–379. 10.1016/j.jmpt.2007.04.001

Zhang, Y., M. Brady, and S. Smith. 2001. “Segmentation of Brain MR Images Through a Hidden Markov Random Field Model and the Expectation-Maximization Algorithm.” IEEE Transactions on Medical Imaging 20 (1): 45–57. 10.1109/42.906424.

